# Thrombospondin module 1 domain (TSP1) of the matricellular protein CCN3 shows an atypical disulfide pattern and incomplete CWR layers

**DOI:** 10.1101/779553

**Authors:** Emma-Ruoqi Xu, Aleix Lafita, Alex Bateman, Marko Hyvönen

**Affiliations:** Department of Biochemistry, University of Cambridge, 80 Tennis Court Road, Cambridge, CB2 1GA, UK; European Molecular Biology Laboratory, Notkestrasse 85, Hamburg, 22607, Germany; European Molecular Biology Laboratory, European Bioinformatics Institute (EMBL-EBI), Wellcome Genome Campus, Hinxton, Cambridgeshire, CB10 1SD, UK

**Keywords:** TSP1 domain, CCN3, crystal structure, conservation analysis, domain definition

## Abstract

Members of the CCN (Cyr61/CTGF/Nov) family are a group of matricellular regulatory proteins, essential to a wide range of functional pathways in cell signalling. Through interacting with extracellular matrix components and growth factors *via* one of its four domains, the CCN proteins are involved in critical biological processes such as angiogenesis, cell proliferation, bone development, fibrogenesis, and tumorigenesis. We present here the crystal structure of the thrombospondin module 1 (TSP1) domain of CCN3 (previously known as Nov), which shares a similar three-stranded fold with the thrombospondin type 1 repeats of thrombospondin-1 and Spondin-1, but with variations in the disulfide connectivity. Moreover, the CCN3 TSP1 lacks the typical pi-stacked ladder of charged and aromatic residues on one side of the domain, as seen in other TSP1 domains. Using conservation analysis among orthologous domains, we show that a charged cluster in the centre of the domain is the most conserved site and predict it to be a potential functional epitope for heparan sulphate binding. This variant TSP1 domain has also been used to revise the sequence determinants of TSP1 domains and derive improved Pfam sequence profiles for identification of novel TSP1 domains in more than 10,000 proteins across diverse phyla.

**Synopsis:** The first structure of a thrombospondin module 1 domain (TSP1) from a CCN family matricellular protein has been determined by X-ray crystallography. The structure shows a typical three-stranded fold, but with an incomplete pi-stacked structure that is usually found in these domains. The structure reveals highest conservation in the positively charged central segment, which we predict to be a binding site for heparan sulphates. The atypical features of this domain have been used to revise the definition of the TSP1 domains and identify a number of new domains in sequence databases.

## 1. Introduction

The CCN proteins are a family of intriguing matricellular proteins, playing regulatory roles in various cellular signalling processes and a range of critical biological functions. There are six members within this protein family in humans, designated CCN1-6 (Brigstock *et al.*, 2003). The CCN acronym was derived from the three prototypical members: <u>**C**</u>yr61 (cysteine-rich protein 61)/CCN1, <u>**C**</u>TGF (connective tissue growth factor)/CCN2, and <u>**N**</u>ov (Nephroblastoma overexpressed gene)/CCN3 (Bork, 1993). The Wnt-inducible proteins, Wisp1-3, are later defined as the three remaining members (CCN4-6) of the family (Pennica *et al.*, 1998).

The CCN proteins have been shown to be involved in developmental processes such as angiogenesis, osteogenesis, proliferation and differentiation (Kubota & Takigawa, 2007; Katsube *et al.*, 2009; Kawaki *et al.*, 2011; Hara *et al.*, 2016); as well as being responsible for diseased states of inflammation, fibrosis, and various types of cancer (Kular *et al.*, 2011; Riser *et al.*, 2015; Li *et al.*, 2015; Kim *et al.*, 2018). However, the molecular mechanism underlying the functions and regulations of the CCN proteins are poorly understood, mostly due to the large number of ligands that have been reported to interact with the CCN proteins and the variety of signalling pathways they are involved in. While CCN proteins have been shown to activate specific signalling pathways, direct receptors for these proteins have not yet been identified. There is also an increasing amount of evidence that CCN proteins, sometimes referred to as growth factors, affect several signalling pathways *via* direct interactions with cytokines and extracellular matrix components. For instance, CCN2 downregulates bone morphogenetic protein (BMP)-2 and -4-mediated Smad1/5/8 phosphorylation and the activation of mitogen-activated protein kinase (MAPK) pathways, thereby inhibiting embryogenesis and chondrocyte proliferation (Abreu *et al.*, 2002; Maeda *et al.*, 2009). Signalling by other growth factors, such as the transforming growth factor β (TGFβ), fibroblast growth factor (FGF), vascular endothelial growth factor (VEGF), and platelet-derived growth factor (PDGF) have also been shown to be affected by the CCN proteins (Abreu *et al.*, 2002; Inoki *et al.*, 2002; van Roeyen *et al.*, 2008; Nishida *et al.*, 2011; Aoyama *et al.*, 2012). Furthermore, the modulation of the signal transduction by the CCN proteins can also be achieved by association with extracellular matrix components. These include sulphated proteoglycans, fibronectin, decorin, low-density lipoprotein (LDL) receptor-related protein (LRP), Notch, and integrins (Chen *et al.*, 2000; Yoshida & Munakata, 2007; Vial *et al.*, 2011; Gao & Brigstock, 2003; Sakamoto *et al.*, 2002; Tan *et al.*, 2009). Despite this wealth of information, the molecular determinants of the interactions of CCN proteins with other molecules are still to be elucidated.

Another feature contributing to the complexity of their molecular functions is that, like many other extracellular proteins, the CCN proteins are a mosaic of structurally distinct domains. Four discrete cysteine-rich domains, an insulin-like growth factor binding domain (IB), a von Willebrand factor C domain (vWC), a Thrombospondin type 1 repeat (TSP1), and a C-terminal cystine-knot domain (CTCK), make up the primary structure of the CCN proteins (Bork, 1993). A short, variable hinge region separates the N-terminal IB and vWC domains from the C-terminal TSP1 and CTCK domains. The CCN family members are highly conserved in their primary structure, with 31-50% pairwise sequence identity between the six paralogs, except for the absence of the CTCK domain in CCN5 (Brigstock, 2003). The 38 cysteine residues are spread out across the four domains and are nearly invariant, with the exception of CCN6 that lacks four cysteines in its vWC domain. Small-angle X-ray scattering (SAXS) analysis has provided us with the first glimpse of the structural arrangements of the four domains, which shows an extended, non-globular fold, with flexibility between the domains predicted to facilitate simultaneous ligand binding (Holbourn *et al.*, 2011). A number of studies have shown that this hinge is prone to proteolysis and the resulting fragments of CCN proteins have been identified in various tissues (Yang *et al.*, 1998; Perbal *et al.*, 1999; Christine P. Burren *et al.*, 1999; Su *et al.*, 2001; Roestenberg *et al.*, 2004). A recent work shows that cleavage of CCN2 in this hinge is required for CCN2-mediated activation of Akt and the ERK pathway, suggesting that the full-length CCN proteins are latent pro-forms (Kaasbøll *et al.*, 2018).

Despite the wealth of data on CCN proteins’ role in signalling, very little is known of their structure and interactions at molecular level. We have recently published the structure of the vWC domain of CCN3 (Xu *et al.*, 2017), but no other high-resolution structures are known for CCN proteins or their domains. Several structures of IB domains have been determined (PDB IDs: 1h59, 1wqj, 2dsp, 3tjq, 3zxb)(Zeslawski *et al.*, 2001; Siwanowicz *et al.*, 2005; Sitar *et al.*, 2006; Eigenbrot *et al.*, 2012; Trachsel *et al.*, 2012) as well as one structure of CTCK domain (Zhou & Springer, 2014), but given their low sequence similarity with CCN proteins, relatively little functional predictions can be derived from these structures.

Here, we present the first crystal structure of the TSP1 domain from CCN3. TSP1 domains, also previously known as thrombospondin type 1 repeats (TSRs), were initially identified in the human endothelial cell thrombospondin-1 (Lawler & Hynes, 1986) and have turned out to be one of the most common motifs in extracellular proteins with close to 402 domains in 97 different human proteins (according to the Pfam database (El-Gebali *et al.*, 2019) release 32.0 at https://pfam.xfam.org/family/PF00090). This small domain contains approximately 50 amino acid residues, and is characterised by a well conserved pattern of residues containing six cysteines (two of which are variable – further details below in Results), two arginines, and two tryptophans.

It has been reported that TSP1 domains inhibit angiogenesis through interactions with α3β1 and αvβ3 integrins, CD36 on the endothelial cell surface, as well as sequestering VEGF away from its receptors (Dawson *et al.*, 1997; Bornstein & Sage, 2002; Inoki *et al.*, 2002). TSP1 domains also typically bear the glycosaminoglycan (GAG) binding sites, used for mediation of GAG-dependent cell adhesion (Clezardin *et al.*, 1997). Neural guidance receptor Unc5 controls the Latrophilin GPCR-FLRT mediated cell adhesion, where its TSP1 domain is responsible for the octameric complex formation (Jackson *et al.*, 2016). In the CCN proteins, the TSP1 domain of CCN2 has been found to display the most promising regenerative effect on chondrocytes and osteoarthritis, compared to the other individual domains and the full-length CCN2 (Abd El Kader *et al.*, 2014). By solving the first structure of a TSP1 domain in the CCN family, we provide the first insights into the possible molecular functions of the CCN proteins.

## 2. Materials and methods

### 2.1. Cloning and expression of CCN3 TSP1

The expression construct of rat CCN3 TSP1 domain (residues 195-249, Uniprot: Q9QZQ5) was amplified by PCR using overlapping oligonucleotides (forward – TATATCCATGGATTCTAGTATCAACTGCATTGAGCAG, reverse – TATATAAGCTTATTCCCCAGGCTCTTGCTCACAAGG) from cDNA (kind gift from Dr. Paul Kemp) and cloned into pHAT4 vector (Peränen *et al.*, 1996), which contains an N-terminal His_6_-tag followed by a TEV protease cleavage site. For protein expression, the construct was transformed into BL21(DE3) *E. coli* competent cells and grown on LB-agar plates containing 100 µg/ml of ampicillin overnight. Resulting colonies were cultured in 2-YT medium with 100 µg/ml ampicillin at 37°C under agitation, until cells reached OD_600nm_ of 0.8-1.0. Protein expression was induced by 400 µM IPTG for 4h at 37°C. Cells were pelleted by centrifugation, resuspended in ddH_2_O, and stored at -20°C.

### 2.2. Protein refolding and purification

The CCN3 TSP1 domain was expressed insolubly in inclusion bodies and subsequently subjected to refolding to regain its native conformation. Harvested cells were first lysed using the Emulsiflex C5 homogeniser in lysis buffer (50 mM Tris-HCl pH 8.0, 2 mM EDTA, 10 mM DTT) with addition of 0.5% (v/v) Ralufon DM detergent. The lysate was incubated with 10 µg/ml DNase I and 4 mM MgCl_2_ for 20 min at room temperature. Inclusion bodies were separated upon centrifugation and washed twice by homogenisation in the lysis buffer containing either 0.5 % Ralufon DM or 1 M NaCl, and finally once with lysis buffer only. Denaturation was achieved by resuspension in 6 M guanidine hydrochloride, 50 mM Tris-HCl pH 8.0, 5 mM EDTA, and 25 mM Tris(2-carboxyethyl) phosphine. The denatured protein was clarified by centrifugation, buffer exchanged to 6 M urea, 20 mM HCl, and adjusted to 1 mg/ml. Refolding was performed by 1:10 rapid dilution into 100 mM Tris-HCl pH 8.5, 100 mM ethanolamine pH 8.5, 1 M pyridinium propyl sulfobetaine, 2 mM cysteine, 0.2 mM cystine, and left for 7 days at 4°C. Refolded protein was purified first by Ni-NTA affinity chromatography, followed by cleavage of the His-tag by TEV protease, and finally by reversed phase chromatography (ACE^®^ 5 C8-300). Purified protein was lyophilised and resuspended in ddH_2_O. MALDI mass spectrometry analysis was used to confirm the molecular weight and the formation of disulfide bonds.

### 2.3. Crystallisation

Purified CCN3 TSP1 domain at a concentration of 17.6 mg/ml was subjected to crystallisation experiments in 96-well plates in sitting drops consisting of 100 nl of protein and 100 nl of crystallisation solution using a number of commercial crystallisation screens. Initial crystal hits were improved by streak seeding using a rabbit whisker, and larger crystals subsequently appeared in 3.0 M NaCl, 0.1 M Tris pH 8.0, in 1µl + 1 µl hanging drops. For experimental phasing, derivative crystals were produced by soaking the native crystals in 6.7 mM K_2_PtCl_4_ in mother liquor overnight. The crystals were cryo-protected in 25% (v/v) glycerol and cryo-cooled using liquid nitrogen.

### 2.4. Data collection, structure determination and refinement

X-ray diffraction data of CCN3 TSP1 were collected at Europe Synchrotron Radiation Facility (ESRF), beamline ID14-4, using an ADSC Q315r CCD based X-ray detector (ADSC, CA), *via* remote control from Cambridge, UK. Multi-wavelength anomalous dispersion (MAD) phasing experiment was performed for the K_2_PtCl_4_ soaked CCN3 TSP1 crystals. Two datasets were recorded for these derivative crystals, at a wavelength of 1.0717 Å for peak anomalous signals, and at 1.0721 Å for inflection point. High resolution diffraction data for the native crystals were obtained at wavelength 0.93 Å. Data were indexed and integrated using *iMOSFLM 1.0.7* (Battye *et al.*, 2011), and scaled using *Aimless 1.1* (Evans & Murshudov, 2013) in the *CCP4 suite 6.3.0* (Winn *et al.*, 2011). *AutoSHARP 2.8.2* (Vonrhein *et al.*, 2007) was used for MAD phasing. The resulting structure was used as the search model for molecular replacement by *Phaser 2.5.1* (McCoy *et al.*, 2007) for the native dataset. Refinement was performed using *Refmac 5.5* (Vagin *et al.*, 2004) and *phenix.refine* (Adams *et al.*, 2010). *Coot 0.7* (Emsley & Cowtan, 2004) was used for model building and validation. Statistics of data collection and refinement are shown in Table 1. The coordinates and structure factors have been deposited in the Protein Data Bank with accession code 6RK1.

**Table 1.** Data collection and refinement statistics. Values for the outer shell are given in parentheses.

|  |  | K<sub>2</sub>PtCl<sub>4</sub> derivative |  |
| --- | --- | --- | --- |
|  | Native | Peak | Inflection point |
| **Data collection** |  |  |  |
| Temperature (K) | 100 | 100 | 100 |
| Wavelength (Å) | 0.9300 | 1.0717 | 1.0721 |
| Space group | P3<sub>1</sub>21 | P3<sub>1</sub>21 | P3<sub>1</sub>21 |
| Cell dimensions |  |  |  |
| *a*, *b*, *c* (Å) | 52.86, 52.86, 102.4 | 52.54, 52.54, 102.9 | 52.56, 52.56, 102.9 |
| α, β, γ (°) | 90.0, 90.0, 120.0 | 90.0, 90.0, 120.0 | 90.0, 90.0, 120.0 |
| Resolution (Å) | 34.13-1.63<br>(1.66-1.63) | 51.43-2.33<br>(2.42-2.33) | 51.43-2.33<br>(2.42-2.33) |
| *R*<sub>merge</sub><sup>b</sup> | 0.082 (0.894) | 0.135 (0.986) | 0.137 (0.768) |
| <*I*> / σ<*I*> | 13.0 (2.3) | 17.5 (3.5) | 217.3 (9.4) |
| Number of reflections | 162469 (8571) | 151309 (13046) | 129980 (10246) |
| Unique reflections | 21384 (1114) | 7506 (771) | 7505 (770) |
| Multiplicity | 7.6 (7.7) | 20.2 (16.9) | 17.3 (13.3) |
| Completeness (%) | 100.0 (100.0) | 100.0 (100.0) | 100.0 (100.0) |
| **Refinement** |  |  |  |
| Resolution (Å) | 34.13-1.63<br>(1.70-1.63) |  |  |
| No. unique reflections | 21339 |  |  |
| *R*<sub>work</sub> / *R*<sub>free</sub> | 0.162 / 0.184<br>(0.214 / 0.232) |  |  |
| No. atoms | 898 |  |  |
| Protein | 743 |  |  |
| Ligand/ion | 15 |  |  |
| Water | 140 |  |  |
| *B*-factors(Å<sup>2</sup>) |  |  |  |
| Protein | 25.50 |  |  |
| Ligand/ion | 25.97 |  |  |
| Water | 39.18 |  |  |
| R.m.s. deviations |  |  |  |
| Bond lengths (Å) | 0.005 |  |  |
| Bond angles (°) | 0.751 |  |  |

### 2.5. Conservation analysis

The ConSurf server (https://consurf.tau.ac.il) was used to evaluate the degree of evolutionary conservation of each amino acid position in the CCN3 TSP1 domain. Protein sequences of TSP1 domain in different CCN family members across a range of species were extracted from the *Ensembl* database (http://www.ensembl.org/index.html, version 76) (Aken *et al.*, 2016), and aligned and scored for position-specific conservation by the ConSurf Server (Ashkenazy *et al.*, 2016).

### 2.6. Sequence Similarity Network construction

A sequence similarity network was constructed using the domain sequences from the four Pfam families of the TSP1 clan (CL0692): TSP_1 (PF00090), TSP1_spondin (PF19028), TSP1_ADAMTS (PF19030) and TSP1_CCN (PF19???). One thousand sequences from each family were randomly selected and used for an all-against-all BLAST search. All pairwise hits with a score above 35 bits were further loaded into Cytoscape (Shannon *et al.*, 2003) to display the network using the default “Perfuse Force Directed Layout” method. In order to highlight the structural coverage of the families in the network, domain sequences in the ECOD database (Cheng *et al.*, 2014) version 248 (23/08/2019) matching any of the TSP1 Pfam family models were included using the same procedure described here.

## 3. Results and discussion

### 3.1. The crystal structure of CCN3 TSP1 domain

The structure of CCN3 TSP1 domain was determined by MAD phasing using K_2_PtCl_4_ derivative crystals (Table 1 and Fig. 1*a*). The three main heavy atom sites are from three platinum atoms bound to sulfur atoms in methionine residues. One Pt bound to methionine 231 in chain A out of two in the asymmetric unit with an occupancy of 1, and two Pt bound to two split conformations of the same M231 in the other chain with occupancies of 0.6 and 0.4, respectively. Two additional sites with low occupancies of 0.2-0.3 are from Pt bound to cysteines when dissociated from disulfide bonds as a result of radiation damage. The structure from MAD phasing was further refined to a resolution of 1.63 Å using the native dataset (Table 1 and Fig. 1*b* and *c*). The two molecules in the asymmetric unit are nearly identical, with an RMSD of 0.345 Å for 44 Cα atoms.

**Figure 1.**
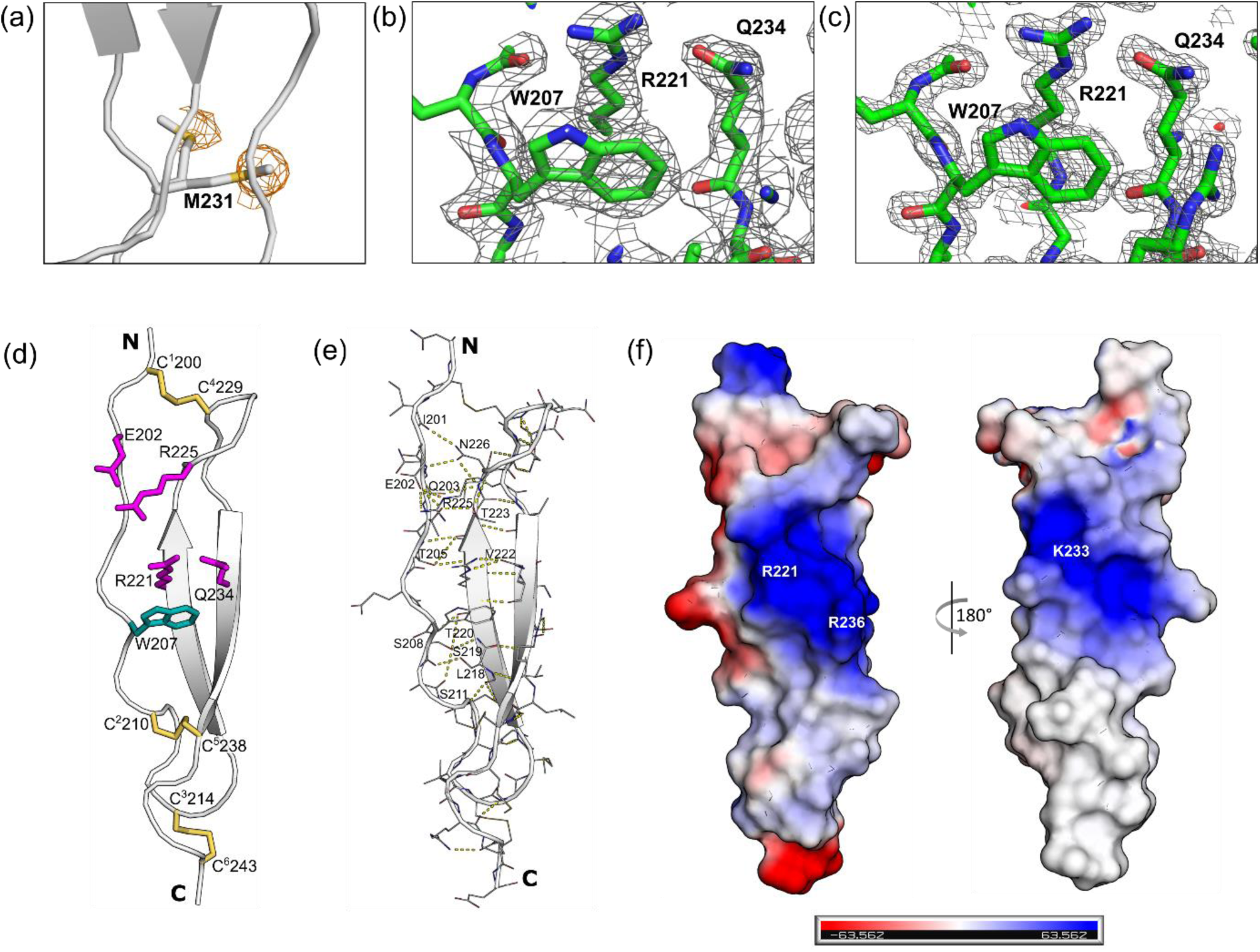
Structure of CCN3 TSP1 domain. (a) Anomalous difference density of Pt sites shown as mesh contoured at 4 σ, with the side chains of M231 bound to Pt atoms shown as sticks, and the protein chain B as ribbon diagrams. (b) Representative 2Fo-Fc electron density map from MAD phasing at resolution of 2.90 Å, contoured at 2 σ. (c) Final 2Fo-Fc electron density map refined to 1.63 Å, contoured at 2 σ. In both (b) and (c) the final, fully refined structure is shown as sticks. (d) Structure of CCN3 TSP1 domain as ribbon diagrams with residues involved in the ‘CWR layers’ shown as sticks. (e) Intra-strand hydrogen bonding in the domain. (f) Electrostatic surface of the TSP1 domain from two orientations.

The CCN3 TSP1 structure exhibits an elongated fold consisting of three antiparallel strands, placing the N- and C-termini at the opposite ends of the domain. This small domain is stabilised by the three disulfide bonds from its six cysteines that are distributed all along the sequence. The top disulfide (when the domain is viewed with its N-terminus pointing up) is formed between C^1^200-C^4^229 (superscripted number refers to the sequential position of the cysteine in the domain) between strands I and III, C^2^210-C^5^238 links strand I to the end of the β-sheet in strand III in the middle of the domain, and third disulfide between C^3^214-C^6^246 links the turn between strands I and II with the very C-terminus of the domain (Fig. 1*d*).

Strand I (N199-C214) is more irregular and rippled, while strands II (G218-L223) and III (Q232-E237) form a regular anti-parallel β-sheet (Fig. 1*d*). In addition to secondary structure-defining interactions between strands II and III, the structure is stabilised by hydrogen bonds formed between the irregular strand I and strand II. These include main chain–main chain atoms of Q203 (O)-N226 (N), T205 (N)-V222 (O), S208 (N)-T220 (O), and S211 (N)-L218 (O), a pair of side chain–side chain H-bonds of E202 O_δ_ with the N_η_ and N_ε_ of R225, and a few other main chain–side chain H-bonds between strand I and II (Fig. 1*e*).

### 3.2. Conservation analysis of TSP1 in CCN proteins

The surface of CCN3 TSP1 domain shows now significant cavities, as potential binding sites for ligands. Projection of electrostatic potential on the surface shows a strongly positively charged zone around the centre of the domain. TSP1 domains are known to bind heparan sulphates (HS) (Guo *et al.*, 2006) and this patch could form a part of a HS binding site. In the absence of mutagenesis or other data on functional sites on TSP1 domain, we turned to the analysis of the evolutionary conservation of the domain. We took all available CCN family proteins from Ensembl genome database and aligned these. This alignment across higher eukaryotes was mapped on to the CCN3 TSP1 structure using ConSurf server by colouring according to conservation scores (Fig. 2). In addition to the almost invariant cysteines, the highest conservation mapped to the central part of the domain, with residues W207, S219, R221, Q234, and R236 showing 100% conservation in all CCN family TSP1 domains. These residues are localised in the positively charged cluster and point to the “front” of the domain, as shown in Figure 1. As they are part of the charged/aromatic spine of the domain, it is impossible to say whether they are conserved for functional or structural reasons, or both.

**Figure 2.**
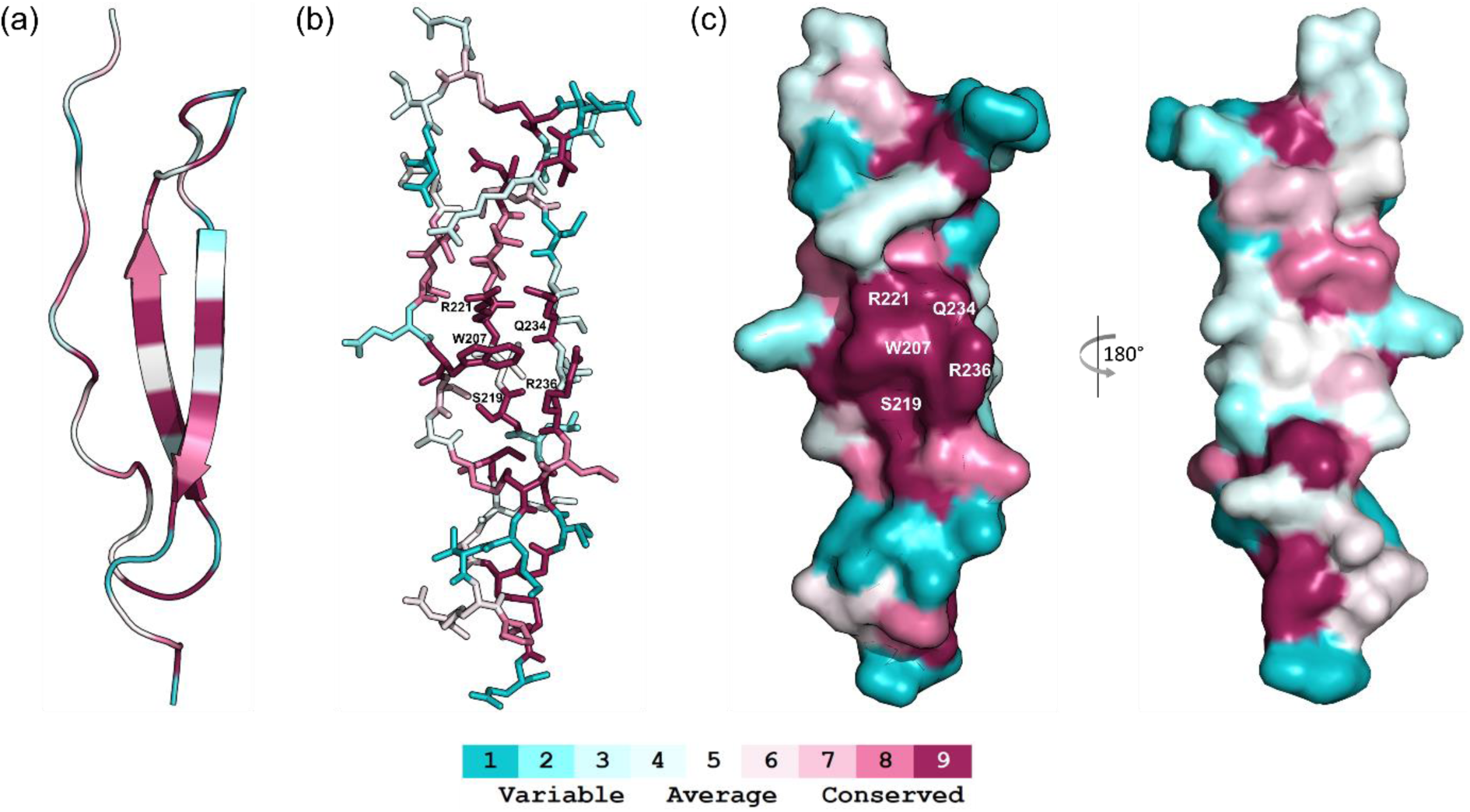
Structure of CCN3 TSP1 reflecting evolutionary conservation. From left to right, are ribbon (a), sticks (b), and surface (c, front and back views) representations, coloured by projecting the conservation scores of the residues (sequence alignment in supplementary Fig. S1) onto the structures. Residues clustered in the most conserved surface patch are labelled.

### 3.3. Similarity and diversity in TSP1 domains

To analyse the structure further, we collected all other TSP1 domain structures from the PDB. Firstly, we used Dali server (http://ekhidna2.biocenter.helsinki.fi/dali/, (Holm & Laakso, 2016) to find the closest homologue from the PDB90 set which contains proteins with maximum 90% pairwise identity. This revealed sporozoite surface protein 2 (PDB 4hqo) as the closest homologue of CCN3 TSP1 domain, but thrombospondin, F-spondin, Complement C6 and C8 proteins were identified in this search as related structures. Further TSP1 structures were taken from the Pfam database listing (family PF00090). List of currently available TSP1 domain structures is shown in supplemental Table S2.

The elongated three-stranded fold is observed in all TSP1 domains. A distinctive feature of the TSP1 domains are the so-called ‘CWR layers’, consisting of the side chain stacking of cystines, tryptophans, and arginines, as described by Tan *et al.* (2002). In the thrombospondin-1 repeat 2/3, an array of tryptophan and arginine residues form multiple π-cation interactions between the aromatic rings and the planar cationic guanidinium groups; these arginines are often found paired with large polar residues forming side-by-side hydrogen bondings across the strands. Together with the three “C” layers of disulfide bridges, these stacked residues form a stabilising spine for this small domain and appear to provide structural rigidity to it in the absence of a hydrophobic core. The first W layer is in some cases replaced by another hydrophobic residue (Leu or Tyr in Spondin-1), but otherwise the CWR-stacked structure is conserved in Spondin-1 repeat 1/4 (Fig. 3a) and structures of other TSP1 domain containing proteins, thrombospondin-repeat anonymous protein (TRAP), complement component C6 and C8 (Tossavainen *et al.*, 2006; Lovelace *et al.*, 2011; Song *et al.*, 2012; Aleshin *et al.*, 2012). By contrast, in the CCN3 TSP1 domain the three R layers and three C layers are conserved, but only the third W layer is present (Fig. 3). The W layers in all TSP1 domains are located in strand I, with their aromatic side chains extending towards the centre of the structure. The R layers in thrombospondin-1 and Spondin-1 TSP1 domains are all formed between strands II and III. In CCN3 TSP1, the top R layer is formed between strands I and II instead. The consequence of the missing top W layer (and the absence of another hydrophobic residue that could replace it), is that the CCN3 TSP1 domain is more open, with strand I more separate from the rest of the structure compared with domains that pull the N-terminus towards the core of the domain by the W layer interactions (Fig. 3).

**Figure 3.**
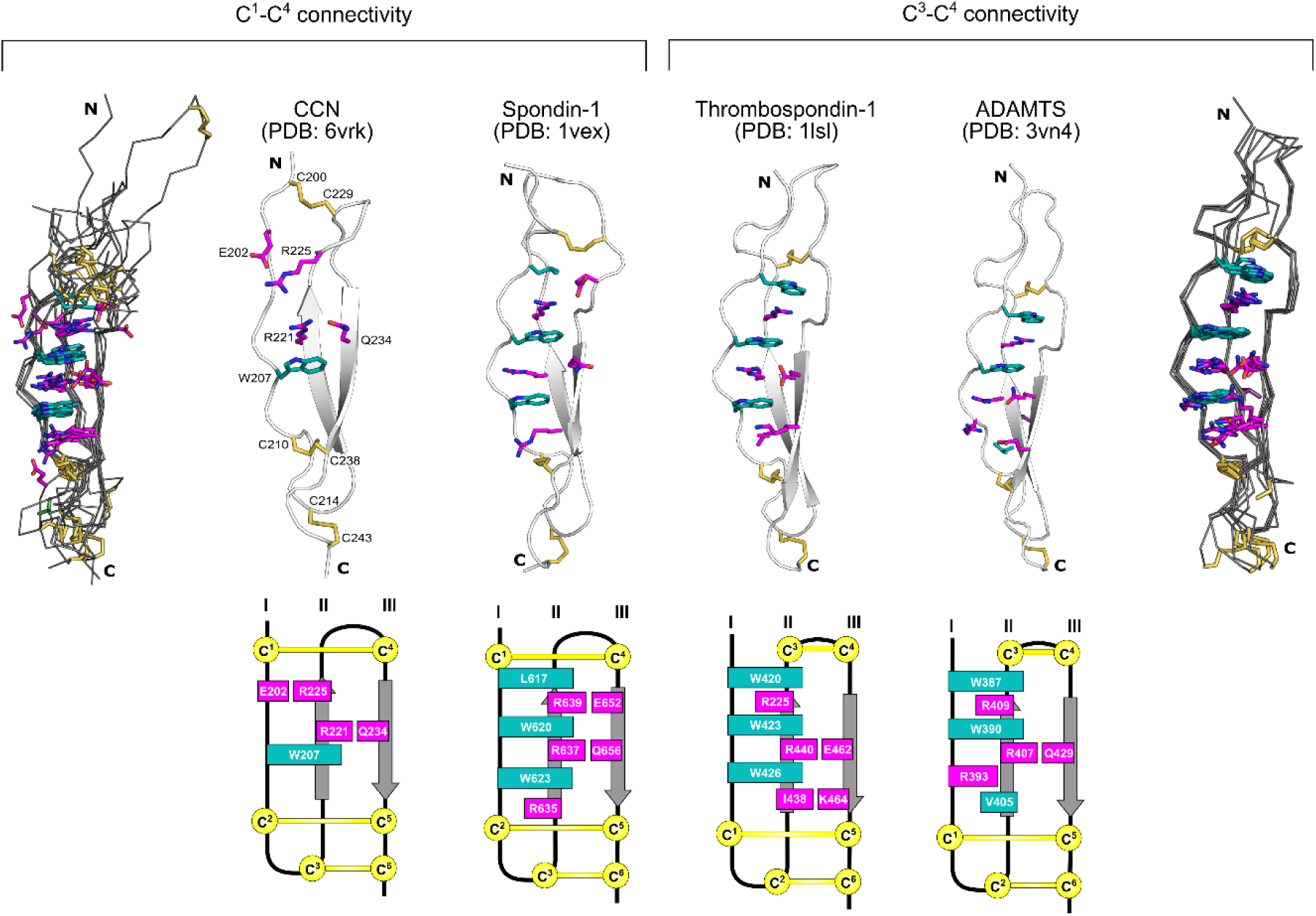
Structural comparisons of TSP1 domains. (a) Ribbon diagrams of TSP1 domains from CCN3 (PDB 6vrk), Spondin-1 (PDB 1vex, repeat 4), thrombospondin 1 (PDB 1lsl, repeat 3) and ADAMTS13 (PDB 3vn4) are shown, with the CWR layers shown in sticks (C-layer in yellow, W layer in teal and in R layer in magenta, including other polar residues that interact with the Arg in the same layer). Schematic topologies of the same domains are shown under the structures, using the same colouring scheme. Superimpositions of Cα coordinates of all TSP1 domains with either C^1^-C^4^ (far left) or C^3^-C^4^ (far right) connectivity are also shown.

Another variable feature among the TSP1 domains is the disulfide bond pattern. The three C layers comprise of one layer at the very top of the structure (when viewed with N-terminus at the top), and two consecutive layers at the bottom, alternated with W and R layers. The bottom two C layers are conserved among all TSP1 domains, formed between strands I and III whereas the top C layer varies in its position and connectivity. In CCNs and Spondins, C^4^ is located at the top of strand III and disulfide bonded to C^1^ at the very N-terminus of strand I. In thrombospondin, C^4^ forms a disulfide with C^3^ (which is missing from CCN-like domains) in the middle of the sequence, at the top of strand II. The differences in the disulfide connectivity in the first C layer at the top of the domain and the lack of the first W layer results in larger differences in the structure of CCN3 TSP1 domain compared with other similar domains. The central W layer in CCN3 ensures that the core of the domain aligns well with other TSP1s. Typical to a large family of disulfide-rich domains, there are always more subtle variations to the connectivity. For example, circumsporozoite protein TSP1 domain (PDB 3vdl and 6b0s) (Doud *et al.*, 2012; Scally *et al.*, 2018) lack the top disulfide and has a long helices containing insertion in loop II-IIII, whereas Micronemal protein 2 (PDB 4okr) (Song & Springer, 2014) contains also a long insertion in loop II-III with an additional pair of disulfide linked cysteines (supplemental Fig. S2).

Overall, the “canonical” TSP1 domains with C^3^-C^4^ connectivity are more structurally conserved with very well defined layered structure, whereas domains with C^1^-C^4^ connectivity have more variable structures and are difficult to align unambiguously.

This difference in the top C layer can be used to categorise different TSP1 domains in matricellular proteins. Sequence alignment of selected TSP1 domains with alternative disulfide connectivities shows that while CCN and Spondin proteins share the same C^1^-C^4^ disulfide pattern, ADAMTS (an extracellular protease), UNC5C (receptor for Netrin), Properdin (a plasma protein), together with thrombospondin, form the alternative group (Fig. 4). However, the functional implication of this structural division of disulfide connectivity is as yet unclear.

**Figure 4.**
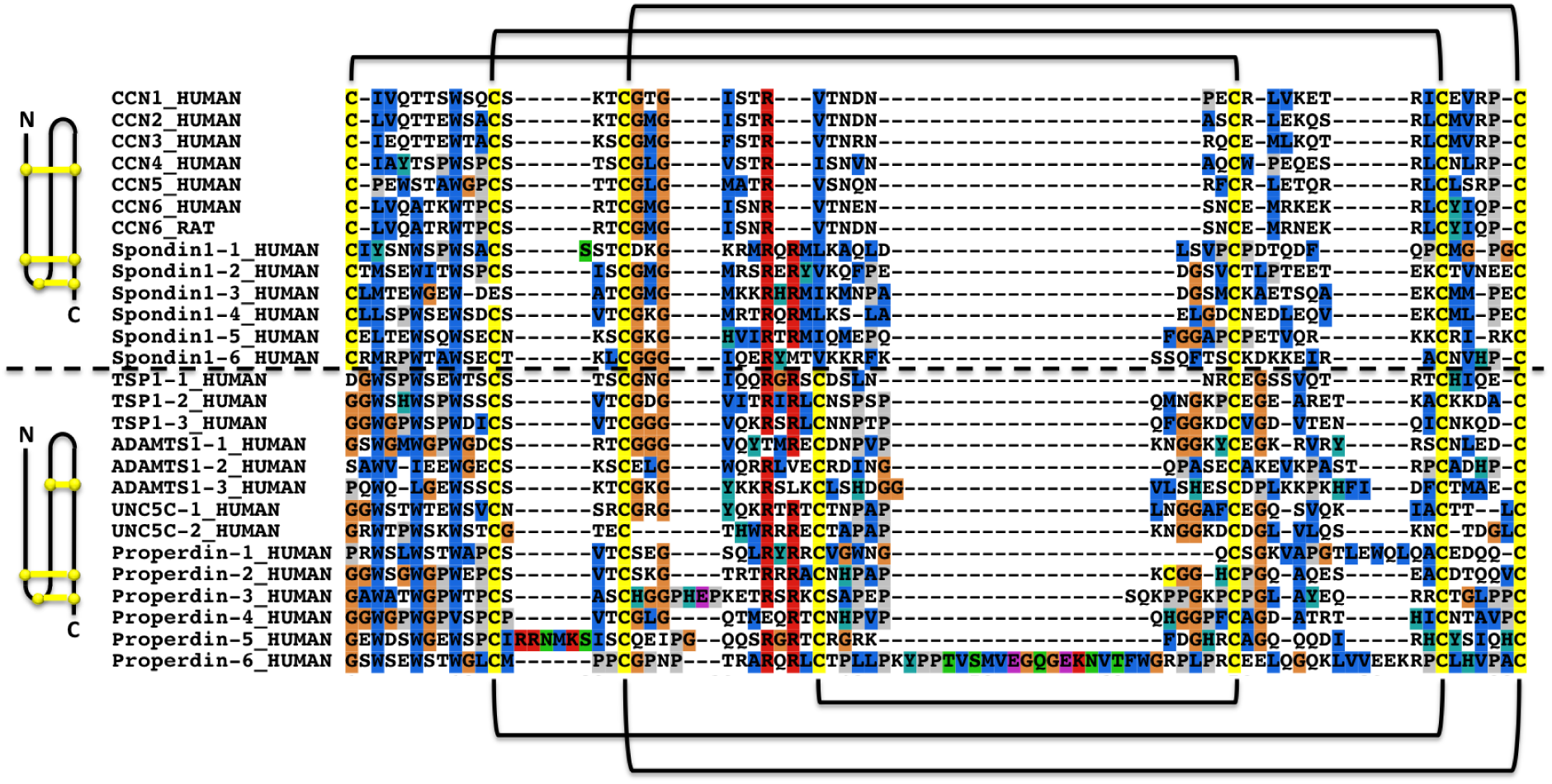
Multiple sequence alignment of the TSP1 domains from CCN1-6 (Uniprot O00622, P29279, P48745, O95388, O76076, O95389), Spondin-1 repeats 1-6 (Uniprot Q9HCB6), thrombospondin-1 repeats 1-3 (TSP1-1 to -3), ADAMTS1 repeats 1-3, UNC5C repeats 1-2, and Properdin repeats 1-6. Dashed line in the middle divided the TSP1 domains into two groups according to their disulfide bond patterns, with schematic representation on the left and connectivity highlighted on the top and bottom of the sequences.

### 3.4. Redefinition of TSP1 domain

The existing Pfam family sequence profile (release 32.0) was entirely built from sequences that had the C^3^-C^4^ connectivity. This meant that matches to C^1^-C^4^ connectivity domains were partial and thus missing the first critical cysteine residue. To remedy this, two new Pfam families were constructed to represent the two subtypes of C^1^-C^4^ connectivity TSP1 domains related to those found in spondins and CCN proteins. These domain families have been deposited in Pfam with accession names TSP1_spondin (PF19028) and TSP1_CCN (PF19???). A new Pfam clan was also built to represent TSP1 domains with accession CL0692 (new TSP1 families and the new clan will be released with version 33.0 of Pfam database). We were further interested to know how common each of these connectivities were across known TSP1 domains. To investigate this, we constructed a Sequence Similarity Network (SSN) of all domains and highlighted the different families. The SSN showed an additional group of sequences that were matching the original TSP1 family but formed their own separated cluster, so a fourth TSP1 Pfam family was built to represent them. This new family (PF19030) was found to correspond to a set of domains from ADAMTS proteins which lack one of the tryptophans, but replace one of the conserved arginines with a hydrophobic residue. Examples of this domain, for which no structures have been experimentally determined yet, can be seen as the second and third domains in ADAMTS1 in Figure 4. The complete SSN displaying the relationships among domains in these three families is shown in Figure 5, along with consensus sequence logo for each of the three families.

**Figure 5.**
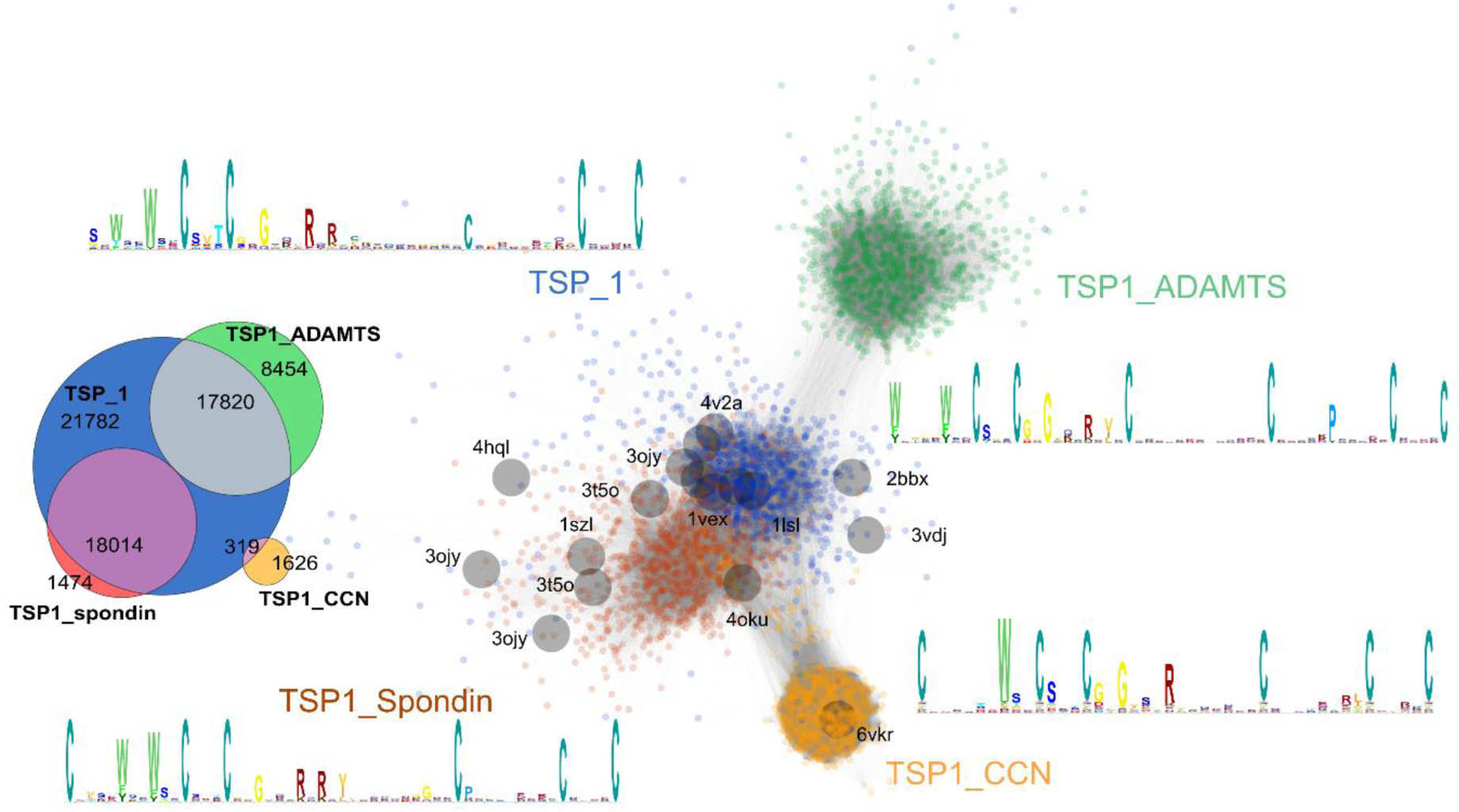
New TSP1 domain families. A Sequence Similarity Network (SSN) of TSP1 domains coloured according to the updated Pfam family definitions. Nodes represent domain sequences and edges represent BLAST hits with a score above 35 bits. Orange nodes belong to the newly defined CCN type TSP1 family (PF19028), maroon nodes correspond to spondin-like TSP1 domains, green nodes to the additional ADAMTS type TSP1 domain family (PF19030), and blue nodes mark the original Pfam TSP_1 family (PF00090). Domains for which experimental structures are known, as derived from the ECOD database, are shown as larger grey nodes and labelled by their PDB accession codes. In the Venn diagram segments of the diagram that do not overlap with the blue circle (original Pfam family PF00090) represent newly identified TSP1 domains. The sequence logos show the conservation within each TSP1 family with the height of the letters correlating with conservation.

Updating the Pfam domain definitions has several important consequences. Firstly, the overall detection of TSP1 domains has increased by 17% from 57,847 to 69,393 (Fig. 5). This includes 13 proteins in SwissProt, four of which are human CCN family members where TSP1 domains were previously not identified by Pfam. Secondly, the improved definitions allow stronger predictions of the disulphide connectivity of Pfam domain matches across all known proteins.

## 4. Discussion

Second X-ray crystallographic structure of a domain from the enigmatic CCN family of matricellular proteins has revealed variant form of TSP1 domain with limited pi-cation ladder typical to these proteins. While functional prediction from this structure of a small domain with limited conservation being difficult, the structure has helped with definition of the larger TSP1 domain population more accurately in Pfam domain database and better annotation of these domains in sequence databases.

Methods for the production of CCN proteins and their fragments in high quality will allow us to use them for further analysis of their molecular functions, identification of interaction partners and biophysical characterisation of these interactions *in vitro*. With significant interest in these proteins as therapeutic targets, in fibrotic conditions in particular, correctly folded proteins will facilitate the development of neutralising antibodies against CCN proteins as well.

## Acknowledgements

We thank Dr Gerhard Fischer for his advice on derivative soaking and MAD phasing. We are grateful for the access to and support at the X-ray crystallographic facility at the Department of Biochemistry. We acknowledge ESRF and their beamline staff for access to beamline ID14-4. The authors declare no conflict of interest.

## Supporting information

**Table S1.** Currently avaialbe TSP1 domain structures in the Protein Data Bank. The last column describes the connectivity of the top disulfide bond, with those that lack it altogether marked as n/a.

| UniProt entry | UniProt residues | PDB ID | PDB residues | Connectivity |
| --- | --- | --- | --- | --- |
| ATS13_HUMAN | 388 - 438 | 3VN4, 3GHN | 388 - 438 | 3-4 |
| CO6_HUMAN | 28 - 78 | 3T5O, 4A5W, 4E0S | 7-57 | 1-4 |
|  | 566 - 612 | 3T5O, 4A5W, 4E0S | 545 - 591 | 3-4 |
|  | 85 - 133 | 3T5O, 4A5W, 4E0S | 64 - 112 | 1-4 |
| CO8A_HUMAN | 540 - 582 | 3OJY | 510 - 552 | 3-4 |
| CO8B_HUMAN | 546 - 590 | 3OJY | 492 - 536 | n/a |
|  | 68 - 116 | 3OJY | 14 - 62 | 1-4 |
| M1V0B0_PLAFA | 329 - 377 | 6B0S | 326 - 374 | n/a |
| O00816_TOXGO | 274 - 331 | 4OKR, 4OKU | 274 - 331 | 1-4 (+) |
| Q7K740_PLAF7 | 326 - 374 | 3VDJ, 3VDK, 3VDL | 326 - 374 | n/a |
| Q9TVF0_PLAVI | 241 - 283 | 4HQL, 4HQN, 4HQO | 241 - 283 | 1-4 |
| SPON1_RAT | 446 - 494 | 1SZL | 446 - 494 | 1-4 |
|  | 618 - 665 | 1VEX | 618 - 665 | 1-4 |
| TRAP_PLAFA | 245 - 288 | 2BBX | 6-49 | 1-4 |
| TSP1_HUMAN | 383 - 428 | 5FOE | 1006 - 1051 | 3-4 |
|  | 439 - 489 | 1LSL, 3R6B | 421 - 471 | 3-4 |
|  | 496 - 546 | 1LSL, 3R6B | 478 - 528 | 3-4 |
| UNC5D_RAT | 254 - 303 | 5FTT | 254 - 303 | 3-4 |
| UNC5A_HUMAN | 240-288 | 4V2A | 240-288 | 3-4 |

**Figure S1.**
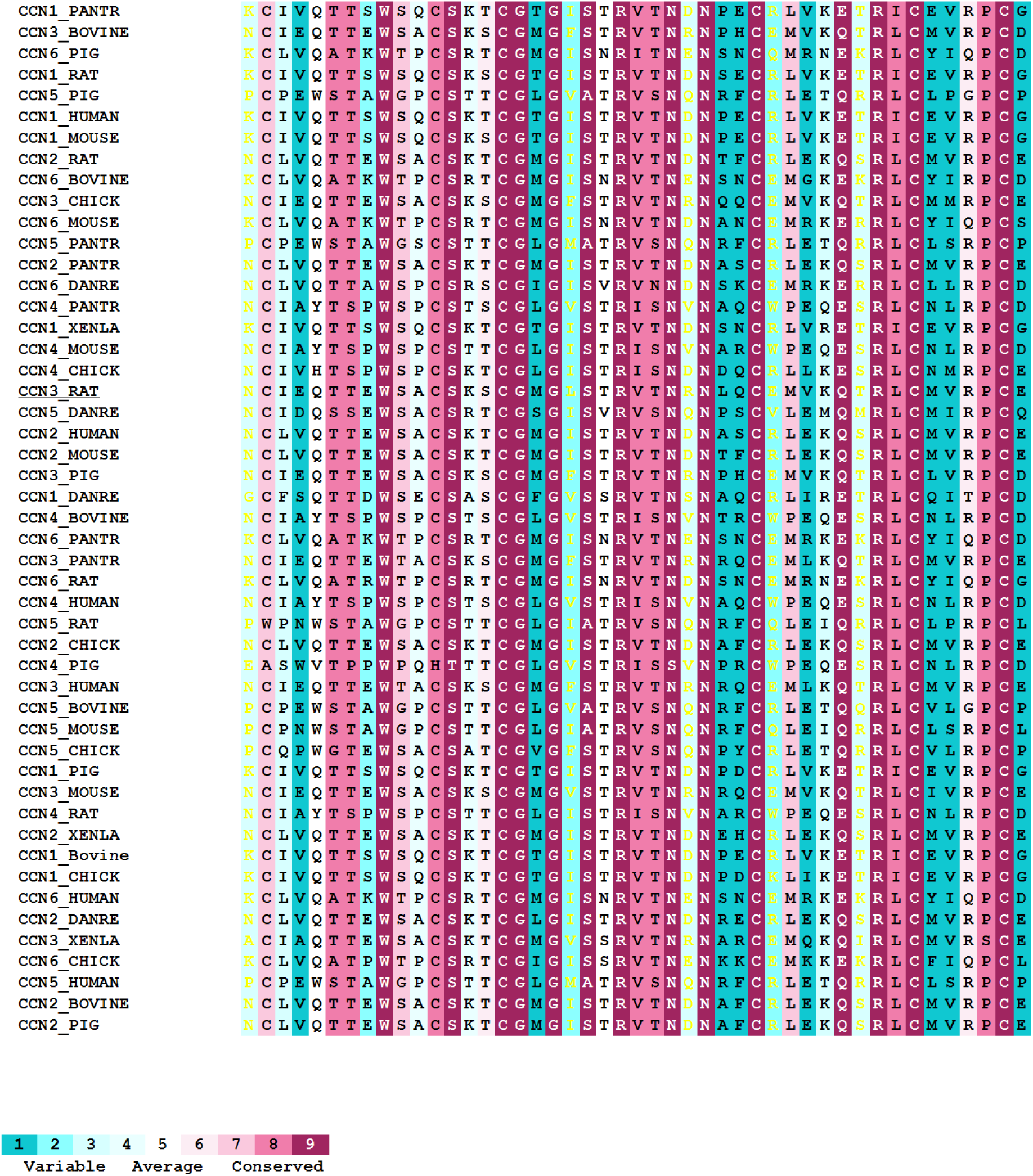
Sequence conservation analysis of the CCN proteins in a range of species generated by the ConSurf Server (https://consurf.tau.ac.il) (Ashkenazy *et al.*, 2010).

**Figure S2.**
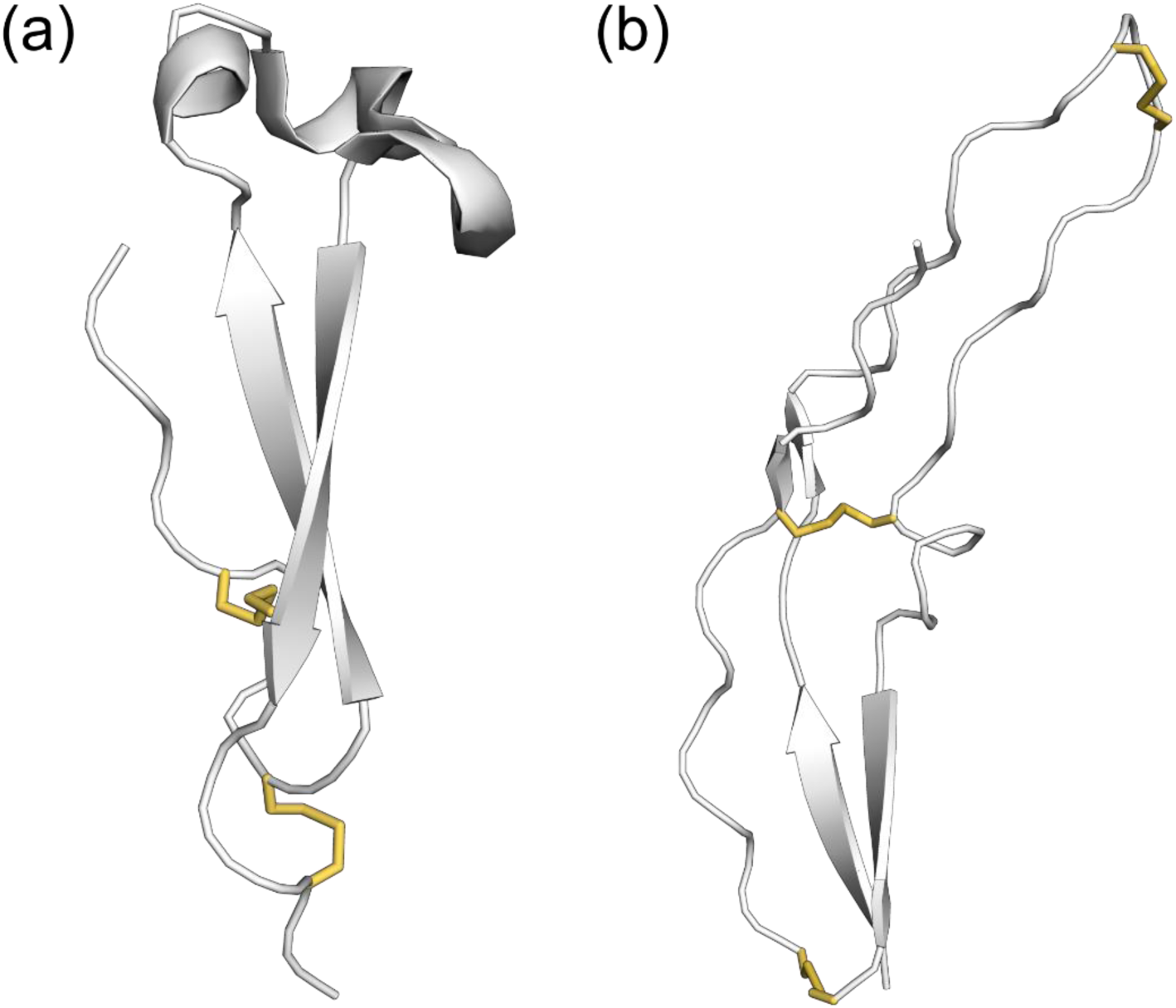
Structure of TSP1 domain from (a) circumsporozoite protein TSP1 domain (PDB 3vdl) and (b) Micronemal protein 2 (PDB 4okr).

